# Genomic and metatranscriptomic analyses reveal an active microbial hydrocarbon cycle in the photic zone

**DOI:** 10.1101/2025.10.24.684404

**Authors:** Kathryn L. Howe, Olivia U. Mason

**Affiliations:** Department of Earth, Ocean and Atmospheric Science, Florida State University, Tallahassee, FL, USA

## Abstract

Natural hydrocarbon seeps and hydrocarbons resulting from human activities are the primary conduits by which oil enters the sea. Marine Cyanobacteria, among the most abundant photosynthetic organisms in the world, produce alkanes, an additional hydrocarbon source to the sea. Alkane production proceeds via metabolism of fatty acid intermediates with a fatty acyl-ACP reductase (FAAR) and an aldehyde-deformylating oxygenase (ADO). These alkanes can be consumed with alkane hydroxylases, including alkane 1-monooxygenase (alkB). The production and consumption of alkanes in the photic zone is termed the short-term hydrocarbon cycle (STHC). Yet, an active STHC has not been substantiated through gene expression analyses. To determine if the STHC is active in the marine photic zone we evaluated over 9000 genomes to which metatranscriptome reads from a site in the Gulf of Mexico were recruited. In these samples, FAAR and ADO expression was dominated by *Prochlorococcus* and to a lesser extent *Synechococcus*. Bacterial alkane consumption via alkB was dominated by Pseudomonadota, SAR324, and Bacteroidota *alkB* gene expression. Additionally, archaeal alkane consumption utilizing this same n-alkane degradation pathway was observed in the transcript data by Thermoplasmatota, classified as Marine Group II and III, the most abundant planktonic archaeal groups. Active production of alkanes in the photic zone would be an important component of marine hydrocarbon cycle and more broadly of carbon cycling. Further, consumption of hydrocarbons, regardless of source, is a fundamentally important ecosystem cleanup service provided by microbes in the ocean.

**Importance:** Hydrocarbons in the long-term hydrocarbon cycle (LTHC) are produced over long-time scales. In comparison, hydrocarbons, specifically n-alkanes, are produced on short time scales by Cyanobacteria as part of basic cellular processes in the short-term hydrocarbon cycle (STHC). Cyanobacterial production of n-alkanes leads to an additional annual input of more than 100 million tons of hydrocarbons to the ocean. Despite the difference in time scale and source, these hydrocarbon cycles are intrinsically linked by microbes that consume n-alkanes using the same pathways for degradation. Here we show that the production and consumption in the STHC is an active cycle in the sunlit ocean, as determined by analysis of genomes and metatranscriptomes. Importantly an active STHC could support a population of alkane degrading bacteria, enabling a rapid response to oil input from a geological reservoir as part of the LTHC, through natural or anthropogenic activities, with implications for bioremediation and ultimately for marine ecosystem health.

## Introduction

Microbes play a pivotal role in marine hydrocarbon degradation, a critical ecosystem service for the remediation of oil spills and natural seepage that is prevalent throughout the Gulf of Mexico (this name is used for historical consistency) and the global ocean. The long-term hydrocarbon cycle (LTHC), the process where oil, millions of years old, is released into the marine environment, either naturally or by drilling and transportation operations, and consumed by microbes, is best exemplified by the bioremediation by planktonic microbes during the 2010 Gulf of Mexico (GOM) Deepwater Horizon (DWH) oil spill.

In contrast, the rapid cycle of cyanobacterial production of hydrocarbons and consumption by microbes occurs over a period of days and is termed the short-term hydrocarbon cycle (STHC). For over 50 years, Cyanobacteria have been known to produce the raw materials for hydrocarbon fuels (Han et al., 1968, 1969; Han & Laboratory, 1968) in the form of medium- and long-chain alkanes (Balaban, 1983; Creton et al., 2010; Schirmer et al., 2010). Roughly 308-771 million tons of these alkanes, specificially pentadecane (C_15_) and heptadecane (C_17_), are produced annually by cyanobacteria, which is ∼1% of the bioavailable dissolved organic carbon in the ocean (Lea-Smith et al., 2015; Love et al., 2021) and ∼100-500 fold more than total oil input into the ocean from all sources (Love et al., 2021; NRC 2003). The functional role of these hydrocarbons in Cyanobacteria was revealed by *in vivo* measurements which showed distortions in cell morphology in knock-out mutants compared to wild type *Synechococcus*; this suggests that these hydrocarbons are required for optimum cell division, size and growth (Lea-Smith et al., 2016). Additionally, they may play a role in response to salinity (Yamamori et al., 2018), cold stress (Berla et al., 2015) and redox balance (Berla et al., 2015).

To date, all cyanobacterial genomes encode hydrocarbon production (Coates et al., 2014), mainly C_15_-C_17_ alka/enes (Lea-Smith et al., 2015) via the FAAR/ADO pathway (Schirmer et al., 2010), which proceeds via metabolism of fatty acid intermediates with a fatty acyl-ACP reductase (FAAR) and an aldehyde-deformylating oxygenase (ADO) (Li et al., 2012). The PKS/OLS pathway is another mode of cyanobacterial alka/ene production, which proceeds via polyketide synthase (PKS) and olefin synthase (OLS) (Mendez-Perez et al., 2011), resulting primarily in nonadecene (C_19_) and similar carbon-length alkanes (Mendez-Perez et al., 2011; Vigneron et al., 2023). However, Coates et al. (2014) found that no cyanobacterial genome encodes both pathways.

Lea-Smith et al. (2015) also showed that alkanes produced by Cyanobacteria were able to support a population of *Alcanivorax borkumensis*, a Gammaproteobacteria considered a paradigm of n-alkane degradation in the marine environment (Schneiker et al., 2006; Yakimov et al., 1998). Love et al. (2021) assembled metagenomes from shipboard incubations with pentadecane and reported ∼20 bacterial genomes encoding the potential for n-alkane degradation, with *Alcanivorax* dominating incubations.

Hydrocarbon degradation by heterotrophic microorganisms was demonstrated more than 100 years ago (Söhngen, 1906) and over 75 genera of hydrocarbon-degrading bacterium have since been recognized (Prince, 2005). In fact, hydrocarbon degradation by marine microorganisms is cited as the reason the ocean is not covered in a layer of oil accumulated from millions of years of natural seepage (Prince, 2005). Recently, Howe et al presented novel MAG and metatranscriptomic read mapping data showing that unculutred Gammaproteobacteria are primarily responsible for degrading aliphatic and aromatic hydrocarbons in the global ocean.

As disccuseed above, hydrocarbon production by marine Cyanobacteria has been shown both in the laboratory by Lea-Smith et al., (2015, 2016) and in shipboard incubations by Love et al. (2021); however, the identity and function of *in situ* producers was not reported, nor was transcriptional activity presented. Further, Love et al. (2021) did not present transcript data on microbial consumption of n-alkanes. Thus, the full cycle of production and consumption in the STHC requires greater resolution on the active players *in situ* through metatranscriptomic read recruitement to genomes. Determining active players and their hydrocarbon degradation pathways could serve to link the STHC and LTHC. To do so, metagenome assembled genomes were mined from the publicly available OceanDNA MAG catalog (Nishimura & Yoshizawa, 2022), which represents a global marine microbial genomic collection of 8,466 species-level clusters that were used in this analysis. Additionally, 920 *Prochlorococcus* and *Synecococcus* genomes in the IMG/Prochlorococcus Portal, a descendant of ProPortal (Kelly et al., 2012), were analyzed herein. Genomes were annoted and metatranscriptomic reads from site GC600, an active hydrocarbon seep in the GOM, were then mapped to these genomes to examine gene expression in microbial hydrocarbon production and consumption pathways as part of the STHC in the GOM and in the global ocean.

## Results

### Hydrocarbon producers, abundance, and genome similarity

Analyses of ProPortal and OceanDNA genomes included gene annotation and read mapping and revealed 604 FAAR/ADO genes that were expressed from 341 genomes (Fig 1, Supp Tables 1 & 2). These genomes were classified as belonging to *Cyanobium*, *Prochlorococcus*, *Synechococcus*, RCC307, AG363K07, or *Phormidesmis* (Fig 1, Supp Tables 1 & 2). Specifically, of the 920 *Prochlorococcus* and *Synecococcus* genomes in ProPortal 1113 FAAR and ADO sequences were identified in 570 microbes. These microbes were sampled from a variety of locations in the global ocean, including epipelagic, pelagic, oceanic, marginal sea, and intertidal zones, and contained 116 cultured representatives (Fig 1, Supp Table 1). Read mapping revealed 575/1113 of these genes were expressed and encoded in 320 microbes which were included in all subsequent analyses (Supp Tables 1 & 2). Of these 320 ProPortal microbes 254 encoded both FAAR and ADO, 28 encoded only FAAR, and 38 encoded only ADO. These same analyses revealed additional genomes encoding and expressing FAAR/ADO genes from the 8,466 prokaryotic species-level clusters in the OceanDNA MAG catalog (Nishimura and Yoshizawa 2022). Specifically, 214 alkane production genes in 108 MAGs in six different phyla were identified, with 29 genes from 21 MAGs from one phylum (Cyanobacteria) that were expressed and included in all other analyses (Fig 1, Supp Table 3). Finally, while some PKS genes were identified in these microbes, no OLS gene annotations were in any of these microbial genomes; therefore, the PKS/OLS pathway was not analyzed any further.

**Fig 1.**
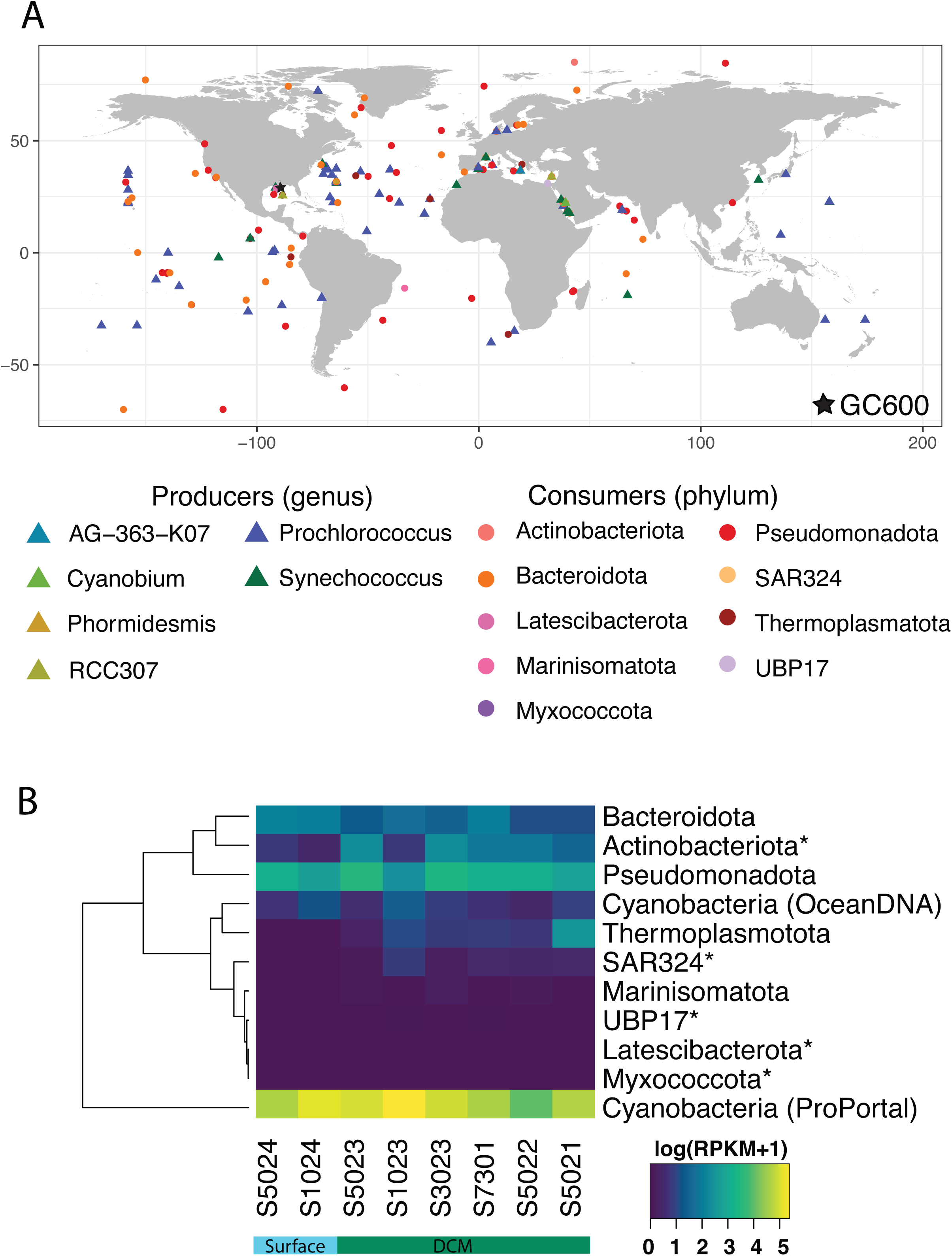
Map showing sample locations for the 504 microbes in this study that expressed at least one alkane production or consumption gene. Color corresponds to clade and shape indicates if the microbe is an alkane producer or consumer. Station location map is also included, with a star marking the GC600 sample site.

Genomes were grouped by phylum revealing that cyanobacterial abundance were the highest of all phyla with summed phylum reads per kilobase per million (RPKM) ranging from 53-217 (Fig 1B), inclusive of hydrocarbon consumers. Specifically, based on read recruitment for individual genomes, *Prochlorococcus* sp. scB243 495P20, assembled from a Bermuda Atlantic Time Series (BATS) site, with a whole genome RPKM of 70 at one depth (Fig 2A) and two *Prochlorococcus* sp. SCGC AC-669 (J8 and L6) assembled from data collected of the coast of Chile (Fig 2A), were the most abundant microbes in the entire dataset. Genome comparison revealed an average nucleotide identity (ANI) among the alkane producers, which were all Cyanobacteria, was > 80% (Supp Fig 1).

**Fig 2.**
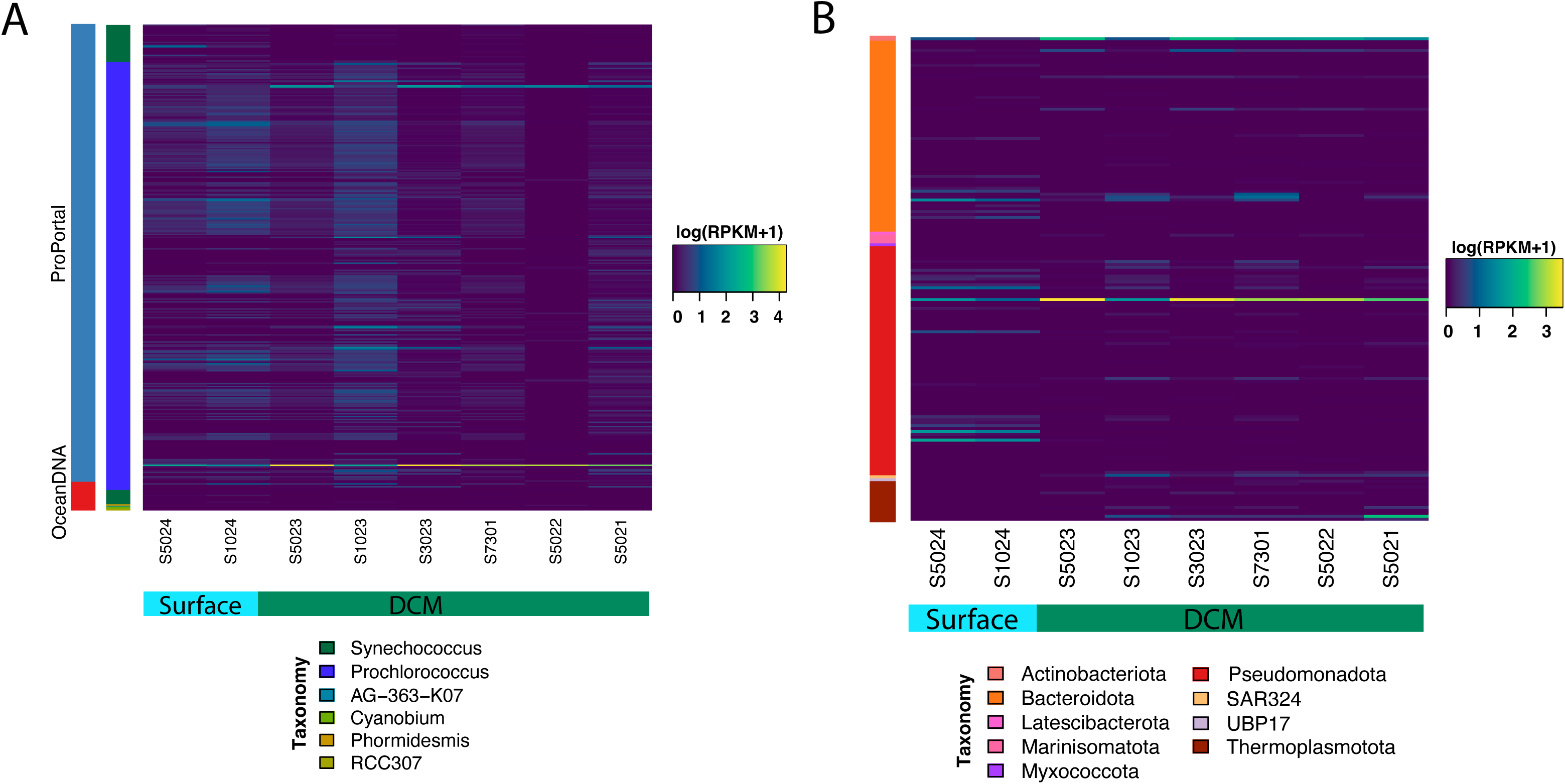
Genome read recruitment for all microbes that express at least one alkane production/consumption gene summed by phylum (A), alkane producers (B), and alkane consumers (C). The * in (A) indicate that phylum is represented by only one MAG. Samples are arranged by increasing depth from left to right. The left color bar in (B) corresponds to the database, while the right color bar indicates genera. The color bar in (C) corresponds to clade.

### Expression of microbial hydrocarbon production genes

ADO and FAAR transcript levels were highest for OceanDNA MAG *Prochlorococcus A* sp. 000635495 (OceanDNA-b16309) assembled from the Red Sea, with an RPKM maximum of 266 and 41, respectively. When summed across all depths RPKM transcript values were 964 for ADO and 177 for FAAR for this organism (Fig 3). Seven other cyanobacterial MAGs from the OceanDNA MAG catalog encoded both FAAR and ADO and recruited reads to one or both genes (Fig 3). Of these, *Prochlorococcus B marinus B* (OceanDNA-b16363) assembled from a GEOTRACES sample in the Mediterranean, had the next highest expression of ADO, with a summed RPKM of 105 across all depths and RPKM > 30 in three deep chlorophyll maximum (DCM) samples, that were defined in Rakowski et al. (2015). FAAR expression for this organism was RPKM maximum of 7.4 at one DCM depth (Fig 3). *Prochlorococcus* sp. MIT9302 had the highest ADO and FAAR gene expression levels for any microbe in ProPortal. The summed expression across all depths was 68 and 13 RPKM, respectively.

**Fig 3.**
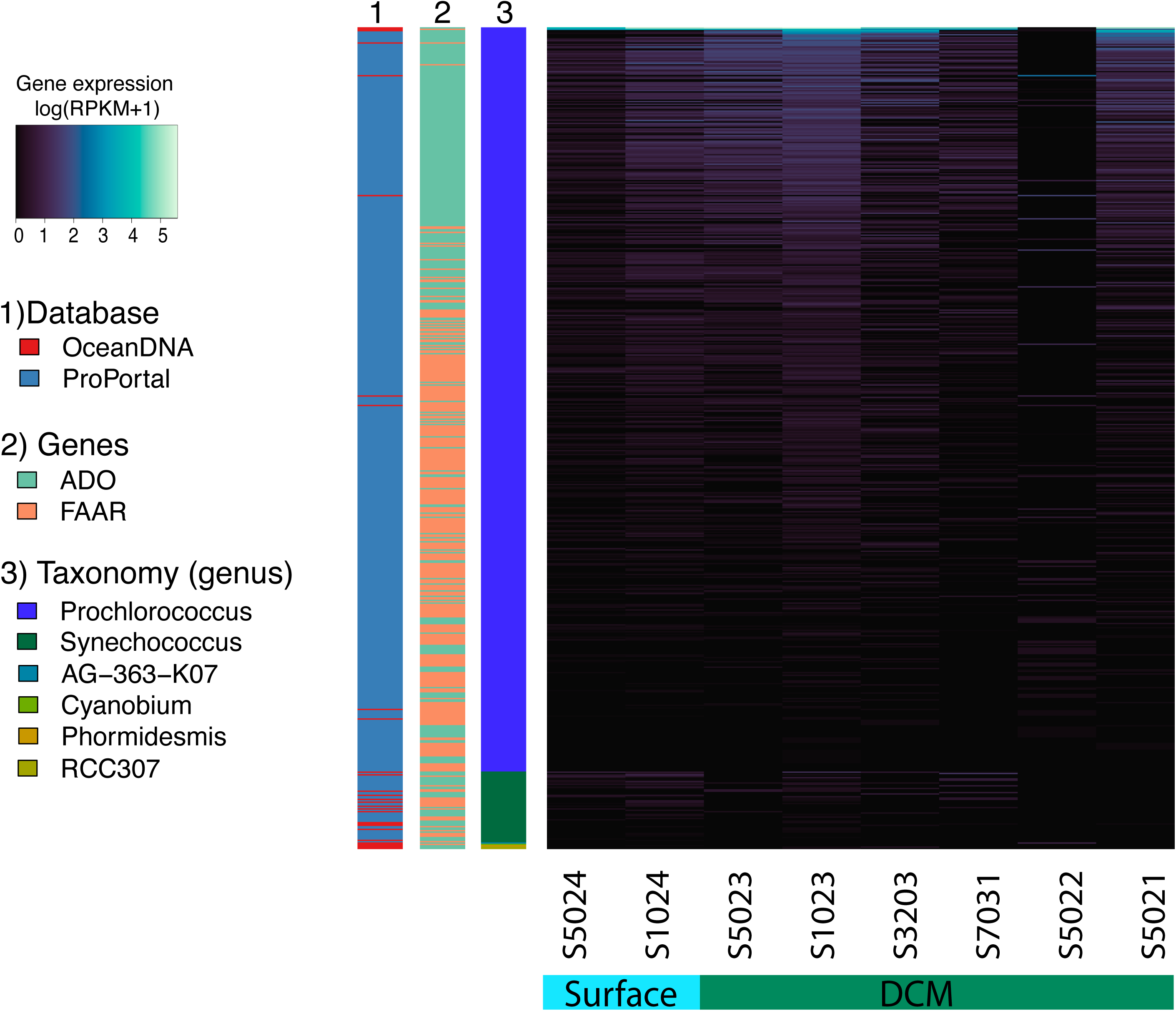
Expression of genes involved in alkane production. Samples are arranged by increasing depth from left to right. As indicated in the key, the first color bar (furthest to the left) corresponds to which database the microbe originated, the next color bar corresponds to taxonomy, and the third relates to which gene is being expressed.

*Synechococcus* genome and gene recruitment was typically much less than that of *Prochlorococcus* genomes (Fig 2A) and genes (Fig 3). *Synechococcus* ADO expression was highest for *Synechococcus* sp. CC9605 with an RPKM maximum of 3, which was the second highest recruitment to any alkane production gene at 82 mbsl by any ProPortal genome (Fig 3). FAAR expression was highest for *Synechococcus* sp. MED850, with an RPKM maximum of 0.94 in a surface sample (Fig 3).

### Hydrocarbon consumers, abundance and, genome similarity

Using the same approach discussed above, consumers were evaluated revealing 2885 *alkB*/CYP153 genes from 2251 MAGs in the OceanDNA MAG catalog spanning 26 phyla, with gene expression observed in nine of these phyla (Fig 4, Supp Table 4). Altogether, 173 alkane hydroxylase (*alkB* and CYP153) genes were expressed in 165 MAGs classified as Actinobacteriota, Bacteroidota, Latescibacterota, Marinisomatota, Myxococcota, Pseudomonadota, SAR324, UBP17 in the Bacteria and Thermoplasmatota in the Archaea (Fig 4, Supp Table 4). Additionally, the metatranscriptomic reads were mapped to the genes annotated as *alkB*/CYP153 presented in Love et al. (2021), but none recruited any reads and therefore were not analyzed further.

**Fig 4.**
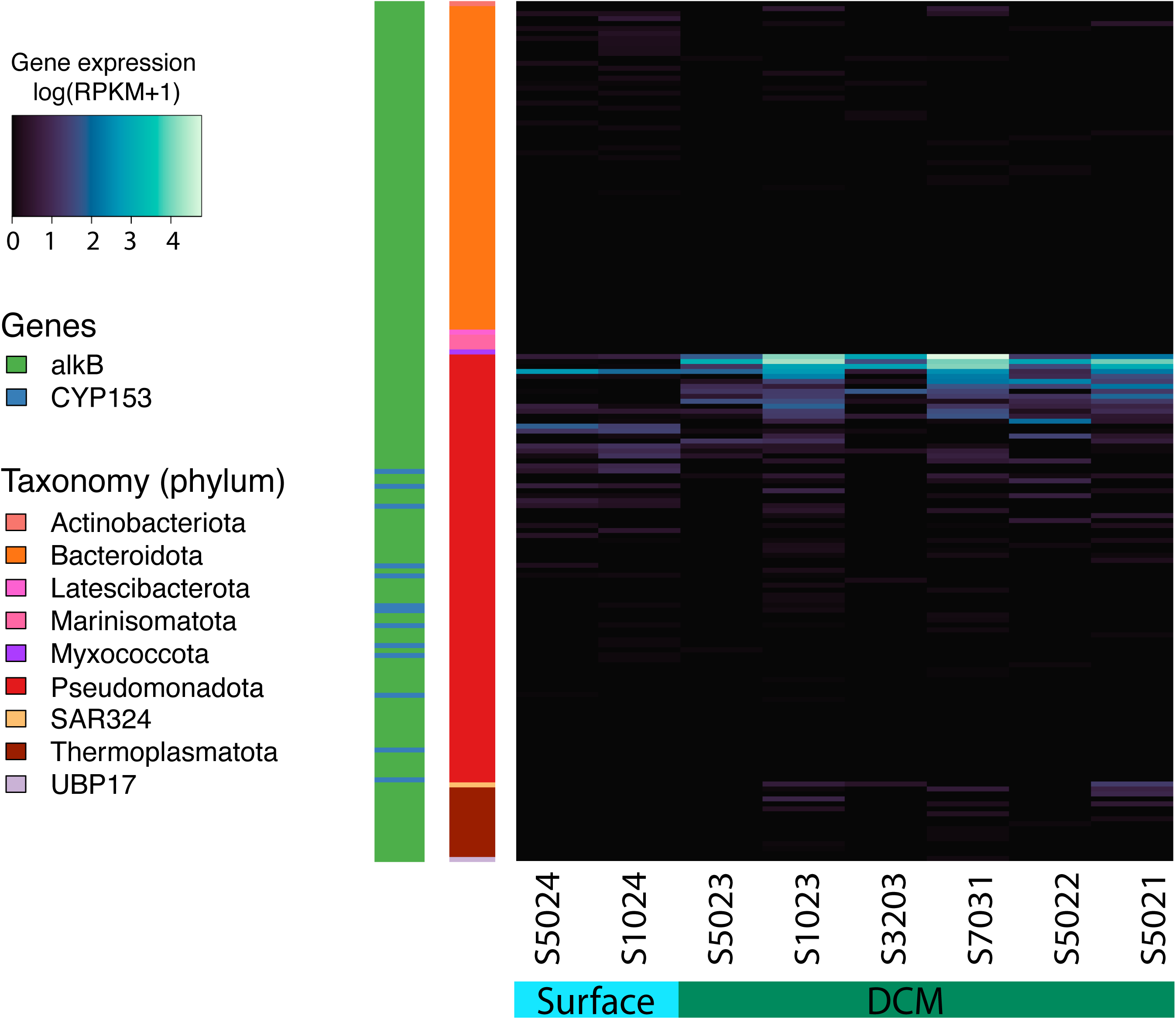
Expression of genes involved in alkane consumption. Samples are arranged by increasing depth from left to right. As indicated in the key, the first color bar (furthest to the left) corresponds to taxonomy, while the other color bar indicates which gene is being expressed.

Genome read recruitment revealed that Pseudomonadota were the most abundant consumers (RPKM range 10-33), followed by Actinobacteriota, Bacteroidota, and Thermoplasmatota (Figs 1B & 2B). The highest genome abundances for individual alkane consumers was an Alphaproteobacteria in the Rhodobacterales (OceanDNA-b30009) with an RPKM range 0.6-9.1, followed by an Actinobacteriota MAG, a member of the archaeal Thermoplasmatota (OceanDNA-a3447) assembled from the Hawaiian Ocean Time-series data, and a Pseudomonadales (OceanDNA-b39211) assembled from Hawaii sea surface (Fig 2B). All remaining alkane consumer genomes had RPKM values of < 5 (Fig 2B).

Given that alkane consumers were identified across several different phyla, ANI was low (Supp Fig 2). For example, Latescibacteriota, Myxococcota, SAR324, UPB17, Marinisomatota, and Actinobacteriota MAGs were < 77% similar to any other MAG. In contrast, within clade similarity was much higher as exemplified by Pseudomonadota, and particularly the Gammaproteobacteria, which were most similar to each other. In this clade the highest similarity (97%) was observed between two *Alcanivorax* genomes, *Alcanivorax* sp. 002354605 (OceanDNA-b36419) and *Alcanivorax* sp. 002354605 (OceanDNA-b36533; Supp Fig 2). Bacteroidota also had high within clade similarity of up to 95% (Supp Fig 2). Finally, the archaeal Thermoplasmatota MAGs in clade similarity was up to an ANI of 93% (Supp Fig 2).

### Expression of microbial hydrocarbon consumption genes

Pseudomonadota were not only the most abundant, but also the most transcriptionally active alkane-consuming clade (Figs 2B & 4). A total of 527 Gammaproteobacteria MAGs encoded 765 alkane hydroxylase genes, with 74 genes from 66 MAGs in 19 orders recruiting reads. These included GCA-002705445, Pseudomonadales, Burkholderiales, UBA4575, Ga0077536, UBA10353, Woeseiales, SAR86, Immundisolibacterales, Nevskiales, and Nitrosococcales (Fig 4). *AlkB* expression was highest in GCA-002705445 microbes, from which 37 *alkB* genes were encoded, with four genes having RPKM values > 10 (Fig 4). In this clade a MAG expressed two copies of its *alkB* gene, both with a summed RPKM of all depths of over 200, with the highest expression in DCM samples (RPKM was 4-177) (Fig 4). Other members of this clade had similar, yet slightly lower *alkB* expression (Fig 4), with Pseudomonadales, the Porticoccaceae, Oleiphilaceae, Spongiibacteraceae families recruiting reads to their *alkB* genes but at lower levels than GCA-002705445 microbes (Fig 4). Interestingly, none of the *alkB* encoded by canonical gammaproteobacterial n-alkane degraders, such as *Bermanella* (3 MAGs), *Cycloclasticus* (8 MAGs), *Oleispira* (8 MAGs), and *Colwellia* (1 MAG), were expressed, except for one *Alcanivorax* (Fig 4).

Altogether, 365 archaeal *alkB* genes were encoded in 361 Thermoplasmatota MAGs, all from Marine Group II (MGII) or Marine Group III (MGIII) in the class Poseidoniia. Of the 365 *alkB* genes, 14 recruited metatranscriptomic reads (Supp Table 3, Fig 4). *Poseidoniaceae MGIIa-L2* sp00249919*5* (OceanDNA-a2828) had the highest *alkB* expression by any archaea (RPKM maximum value of 1.5), followed by *Poseidonia* sp002714305 (OceanDNA-a3447; RPKM maximum of 1.3). *MGIIa-L1* sp8160u (OceanDNA-a2630) recruited metatranscriptomic reads to its *alkB* genes at the most depths (three depths with RPKM range 0.05-1.23) with the remaining archaeal *alkB* genes having minimal expression (Fig 4). Only one archaeal MAG (OceanDNA-a3413) expressed *alkB* in a surface sample (RPKM was 0.02), the rest of the activity was in the DCM (Fig 4).

### CYP153 and secondary alkane consumption genes

Secondary alkane consumption genes, including generic cytochrome P450 (not annotated as an alkane hydroxylase; cytP450) and ferredoxin were expressed by Actinobacterota and Bacteriodota, while only Bacteriodota expressed rubredoxin (Supp Fig 3). Flavin-binding monooxygenase (FMO) were expressed by Bacteroidota, Myxococcota, Planctomycetota, Pseudomonadota, Thermoplasmatota and while ferredoxin reductase genes were encoded, none recruited any reads. Pseudomonadota, specifically the Pseudomonadales (Halieaceae, Alcanivoracaceae, and HTCC2089), was the only clade to express CYP153 genes, but expression levels were low compared to *alkB* expression (Fig 4).

Other genes involved in consumption, such as FMO, was observed in the Pelagibacterales with six MAGs encoding FMO and three expressing it (Supp Fig 3). In addition to the Pelagibacterales MAGs, ten other Alphaproteobacteria expressed FMO, with an RPKM maximum of 8 by *Rhizobales* AG-430-B22 sp003212795 (OceanDNA-b27101) (Supp Fig 3). Other microbes classified as Flavobacteriales SAR86, Ga0077536, and one Myxococcota encoded and expressed FMO genes, some of which also expressed ferredoxin (RPKM max of 3.6), and generic cytP450 (RPKM maximum of 0.04; Fig 4 and Supp Fig 3). Only one Thermoplasmatota MAG expressed FMO (RPKM < 0.04), but no other secondary alkane consumption genes. Actinobacterota and Bacteriodota expressed cytP450 with RPKM maximum of 1.76 (Supp Fig 3). One Planctomycetota classified as Pirellulales (OceanDNA-b22322) expressed FMO with an RPKM <0.02. Only one Bacteriodota MAG classified as Flavobacteriales (OceanDNA-b9761) that expressed rubredoxin expressed any other alkane consumption gene, in this case ferredoxin, yet both were minimally expressed (RPKM < 0.05; Supp Fig 3).

## Discussion

In our study, *Prochlorococcus* were the most abundant alkane producers, with the highest overall FAAR and ADO gene expression levels. In particular, two uncultured *Prochlorococcus* assembled from the Red Sea and from a Bermuda Atlantic Time Series (BATS) sample, recruited the greatest number of reads to their FAAR and ADO genes. In laboratory experiments with cultured *Prochlorococcus*, Lea-Smith et al. (2015) measured hydrocarbon production by *Prochlorococcus* MIT9312 (high-light adapted strain and one of the most numerically abundant *Prochlorococcus* ecotypes) and MIT9313 (low-light adapted strain), both of which were determined to actively express alkane production genes *in situ* in our samples. Lea-Smith et al. (2015) also showed that *Synechococcus* produce n-alkanes, consistent with our findings that members of this group encode and express FAAR and ADO genes *in situ*. Specifically, they showed that *Synechococcus sp.* WH8102, WH7803 and WH7805 produce n-alkanes. All three were found to encode FAAR and ADO in our samples, yet only *Synechococcus sp.* WH8102 expressed these genes *in situ*. Thus, our findings provide the missing information needed to link laboratory observations to *in situ* gene expression for n-alkane production pathways as part of the STHC. Further, inclusion of uncultured microbes from the global ocean that were actively transcribing genes involved in alkane production broaden our understanding of how widespread alkane production is in the marine environment.

In terms of n-alkane consumption, Pseudomonadota were the most abundant and active alkane consumers. In particular, members of the gammaproteobacterial order GCA-002705445 had the highest *alkB* expression levels. This order contains methylotrophs that have been shown to increase in abundance in response to phytoplankton blooms (Francis et al., 2021). Additionally, Porticoccaceae MAGs assembled from a low-oxygen Hawaiian Ocean Time-series sample, the North Sea, and the Mediterranean Sea, expressed *alkB*. Members of this group have also been shown to be associated with phytoplankton and are known to degrade PAHs (Gutierrez et al., 2012; Somee et al., 2022; Yakimov et al., 2022). Other Pseudomonadota, such as Rhodobacteraceae in the Alphaproteobacteria, encoded and expressed *alkB*. Rhodobacteraceae are known to degrade aromatic hydrocarbon compounds (Hu et al., 2017), to colonize plastics (Pinto et al., 2019; Zhang et al., 2022), to be present and active in hydrocarbon-contaminated environments (Lamendella et al., 2014; Mishamandani et al., 2016), and were the predominant bacteria affecting n-alkane concentrations in DWH microcosms presented in Hu et al., (2017). Overall, our findings provide greater resolution on why these microbes that can degrade alkanes, such as those produced by phytoplankton as part of the STHC, may respond to phytoplankton blooms. Further, the ability to degrade hydrocarbons as observed through genome analysis and read mapping extends our understanding on why members of these clades are present in hydrocarbon contaminated environments and what type of hydrocarbons they have the genetic machinery to degrade.

Outside of the Pseudomonadota, Bacteroidota, specifically the Flavobacteriales, Actinobacteriota, Latescibacterota, SAR324, as well as Myxococcota and UBP17 members encoded and expressed *alkB*, with Flavobacteriales being the only group to express all secondary alkane consumption genes, and the only group to express rubredoxin. Members of these clades are known to degrade aromatic and aliphatic hydrocarbons, are often associated with the plastisphere, and some are also known to increase in abundance in response to a phytoplankton bloom (Liu & Liu, 2013; Bunse & Pinhassi, 2017; Dombrowski et al., 2017; Hu et al., 2017; Tu et al., 2020; Love et al., 2021; Somee et al., 2021; Vaksmaa et al., 2021; Somee et al., 2022; Howe et al., 2023, 2024). Thus, our understanding of these microbes is largely that they respond to oil input as part of the LTHC, but our results suggested that, like the Pseudomonadota, they may also play an important role in the consumption side of the STHC.

Archaeal alkane consumption by Thermoplasmatota (class Poseidoniia) using *alkB* was observed in the transcript data. These MAGs were from the Atlantic and Pacific Oceans, as well as the GOM and Mediterranean Sea. Poseidoniia includes MGII and MGIII which are the most abundant planktonic archaeal groups in the surface ocean water column (Rinke et al., 2019) and are frequently observed during or after phytoplankton blooms due to possible syntrophic interactions between these groups (Chen et al., 2022). The limited number of sequenced genomes and lack of cultured representatives of Thermoplasmatota, and specifically MGII and MGII subgroups, leaves a knowledge gap about their metabolic capabilities; however, they have previously been shown to be capable of alkane degradation with *alkB* (Somee et al., 2022; Somee et al., 2021; Rinke et al., 2019) and some of these sequences cluster with gammaproteobacterial *alkB* (Somee et al., 2022). Additionally, analysis of TARA Oceans data by Love et al. (2021) revealed a clade of *alkB*-like monooxygenases belonging to MGII/MGIII that was persistent in all surface and DCM samples of the Northern Atlantic Ocean, suggesting that members of these MGII and MGIII clades may play a role in the STHC. Our study provides new, important information on active transcription of alkane consumption genes in Thermoplasmatota, and MGII and MGIII in particular, that appear to be involved in both the LTHC and STHC.

To evaluate transcriptional activity in the STHC and explore linkages with the LTHC, we selected the Green Canyon 600 (GC600) lease block in the GOM, an area of natural hydrocarbon seeps (Garcia-Pineda et al., 2010) with persistent surface oil slicks (Garcia-Pineda et al., 2010; Krajewski et al., 2018). However, slicks resulting from GC600 seepage are characterized as highly weathered with n-alkanes < C_15_ being largely absent, and C_16_–C_34_ components comprising 3-8% of the n-alkane pool (Ziervogel et al., 2014). Further, at GC600 surface oil slicks are ephemeral with an average residence time of approximately 6.4 hr. The depletion of n-alkanes < C_15_ coupled with the ephemeral nature of oil slicks at this site, may explain why DWH microbes described in Howe et al. (2024) were inactive at GC600, suggesting that while the LTHC and STHC are linked through metabolic pathways for consumption, sustaining a population of hydrocarbon degrading microbes is likely by alkane production in the STHC. To this end, it is estimated that alkanes produced as part of the STHC would represent an additional hydrocarbon source in the global sunlit ocean that could equate to 308-771 million tons of hydrocarbons input annually (Lea-Smith et al., 2015), an amount which exceeds that of hydrocarbons produced by Saudi Arabia (US Department of Energy). This additional substantial input of hydrocarbons outside of the LTHC could sustain populations of alkane degrading microbes, such as those presented herein, and would facilitate rapid microbial consumption of alkanes from the LTHC.

All together, our analysis of over 9000 genomes generated from microbes residing in the global ocean combined with metatranscriptome read mapping, revealed, for the first time, expression of genes involved in both production and consumption as part of the STHC. An active STHC is ecologically consequentional given the vast quanity of hydrocarbons produced by Cyanobacteria that would need to be continuously consumed. Further, these hydrocarbons could support a population of microbes that are poised to degrade hydrocarbons from the LTHC as they transit through the marine water column. Additionally, the abundance and/or expression of *alkB* by marine microbes has been shown to be associated with or contributes to the degradation of some plastics, such as polyethylene and low density polyethylene (Farveen & Narayanan, 2024; Gates & Crook, 2024; Jeon & Kim, 2015; Krueger et al., 2015; Pinto et al., 2022; Salam, 2024; Yoon et al., 1996), suggesting that they play a role in mitigating additional petroleum-derived waste in the form of plastics (Yakimov et al., 2022). Thus, microbes described herein could have an important role in marine health as they are capable of consuming hydrocarbons produced by Cyanobacteria, natural seepage, anthropogenic input, and in the form of plastics. Collectively, this work provides important new insights into marine microbes as ecosystem engineers, actively producing hydrocarbons in the sunlit ocean and subsequenctly consuming hydrocarbons from the deep-sea to the surface ocean, linking the STHC, LTHC and the persistence of marine plastics.

## Materials and Methods

### Sample collection

Eight depths at GC600, an active seep with visible oil at the surface, in the GOM were sampled during an expedition in June 2013 on the *R/V Pelican* (Rakowski et al., 2015). Collection depths spanned the near-surface (< 2 mbsl) to 120 mbsl for these eight depths where the euphotic zone the deep-chlorophyll maximum (DCM) was targeted for collection. The microbial communities at this site were described in Rakowski et al., (2015). These samples from eight depths at GC600 were selected for metatranscriptomic sequencing.

### Microbial sample collection and RNA extractions

From GC600 10 L of seawater was collected from eight depths and filtered with a peristaltic pump. A 2.7 μM Whatman GF/D pre-filter was used and samples were concentrated on 0.22 μM Sterivex filters (EMD Millipore). Sterivex filters were sparged and filled with RNAlater. RNA was extracted directly off of the filter by placing half of the Sterivex filter in a Lysing matrix E (LME) glass/zirconia/silica beads Tube (MP Biomedicals, Santa Ana, CA) using the protocol described in Gillies et al. (2015) which combines phenol:chloroform:isoamyalcohol (25:24:1) and bead beating. RNA was stored at -80°C until purified using QIAGEN (Valencia, CA) AllPrep DNA/RNA Kit.

### Metatranscriptomic sequencing

For metatranscriptomic sequencing only RNA with an RNA integrity number (RIN) (16S/23S rRNA ratio determined with the Agilent TapeStation) ≥ 9.5 (on a scale of 1-10, with 1 being degraded and 10 being undegraded RNA) was selected for sequencing. Using a Ribo-Zero kit (Illumina) rRNA was subtracted from total RNA. Subsequently, mRNA was reverse transcribed to cDNA as described in Mason et al. (2012). RNA (as cDNA) libraries were linker barcoded and pooled for high-throughput Illumina HiSeq 2000 paired-end sequencing at Argonne National Laboratory as described in (Mason et al., 2014).

### Genome analyses and read mapping

The OceanDNA MAG collection contains 52,000 genomes with 8,466 prokaryotic species-level clusters (Nishimura and Yoshizawa 2022) that were analyzed here along with 920 *Prochlorococcus* and *Synecococcus* genomes from the IMG/Prochlorococcus Portal (https://img.jgi.doe.gov/cgi-bin/proportal/main.cgi), hereafter referred to as ProPortal. Average nucleotide identity (ANI) for all genomes was determined using Sourmash with --containment -- ani --ksize 31 (Brown & Irber, 2016). Functional annotations targeting alkane production and consumption were identified using Pfam (Bateman et al., 2004) and KEGG (Kanehisa et al., 2016) annotations through DRAM (Shaffer et al., 2020). Metatranscriptome reads were mapped to the 920 *Prochlorococcus* and *Synecococcus* genomes from ProPortal, using Bowtie2 (Langmead & Salzberg, 2012). The metatranscriptome reads were also mapped to the 8,466 medium to high-quality species-level genomes from the OceanDNA MAG catalog (Nishimura & Yoshizawa, 2022). Lastly, these metatranscriptomic reads were mapped to the genes annotated as alkB/CYP153 in the 16 MAGs presented in Love et al. (2021).

Alkane producers were defined as expressing fatty acyl-ACP reductase (FAAR) and/or aldehyde-deformylating oxygenase (ADO) in at least one sample. Primary alkane consumers were defined as expressing alkane 1-monooxygenase (*alkB*) and/or cytochrome P450 alkane hydroxylase (CYP153) in at least one sample. Although not a criterion for inclusion, secondary alkane-consumption genes including flavin-binding monooxygenase (FMO), generic cytochrome P450 (not alkane hydroxylase; cytP450), rubredoxin, and ferredoxin were included in the analysis to give a more comprehensive view of hydrocarbon degradation capabilities of these microbes. Reads for genome and transcript abundance were normalized by calculating reads per kilobase per million (RPKM) values by following Thrash et al., (2017). Low and minimal gene expression are defined herein as RPKM < 5 and RPKM < 1, respectively.

## Acknowledgements

We thank the captain, crew and science teams aboard the R/V Pelican. This work was supported by a grant from The Gulf of Mexico Research Initiative RFP-VI: Research Grants [OUM Consortium for Simulation of Oil-Microbial Interactions in the Ocean (CSOMIO)].

## Data Availability Statement

Metatranscriptome reads are publicly available in NCBI (PRJNA1346479) and on the Mason server http://mason.eoas.fsu.edu. Publicly available genomes and MAGs were downloaded from https://img.jgi.doe.gov/cgi-bin/proportal/main.cgi (IMG/Prochlorococcus Portal) and https://doi.org/10.6084/m9.figshare.15186039.v1 (OceanDNA).

## Supplemental figures and tables

SF 1. Average nucleotide identity (ANI) of alkane producers.

SF 2. Average nucleotide identity (ANI) of alkane consumers.

SF 3. Heatmap of secondary alkane consumption genes

ST 1: The 320 ProPortal genomes that expressed alkane production genes FAAR and/or ADO.

ST 2: The 21 OceanDNA MAGs that expressed alkane production genes FAAR and/or ADO.

ST 3: The 165 OceanDNA MAGs that expressed primary alkane consumption genes alkB and/or CYP153.

